# Optimization of information transmission through a noisy biochemical pathway explains the choice of reaction rates

**DOI:** 10.1101/2022.07.01.498389

**Authors:** Anush Mahajan, Bhaswar Ghosh

**Affiliations:** International Institute of Information Technology, Hyderabad

## Abstract

Living organisms are required to sense the environment accurately in order to ensure appropriate responses. The cells utilize specific signaling pathways to estimate the signal. The accuracy of estimating the input signal is severely limited by noise stemming from inherent stochasticity of the chemical reactions involved in signaling pathways. Cells employ multiple strategies to improve the accuracy by tuning the reaction rates, for instance amplifying the response, reducing the noise etc.. However, the pathway also consumes energy through incorporating ATP in phosphorylating key signaling proteins involved in the reaction pathways. In many instances, improvements in accuracy elicit extra energetic cost. For example, higher deactivation rate suppresses the basal pathway activity effectively amplifying the dynamic range of the response which leads to improvement in accuracy. However, higher deactivation rate also enhances the energy dissipation rate. Here, we employed a theoretical approach based on non-equilibrium thermodynamics of information to explore the role of accuracy and energetic cost in the performance of a MAPK signaling system. Our study shows that the accuracy-energy trade-off can explain the optimality of the reaction rates of the reaction pathways rather than accuracy alone. Our analysis elucidates the important role of the interplay between accuracy and related energetic cost in evolutionary shaping of the parameter space of signaling pathways.

## Introduction

Every living cell has to accurately sense the external environment in order to ensure appropriate response. The sensing is carried out by cell signal transduction systems which convey the message of external stimulus to the internal machinery of the cell. Various chemical reaction pathways are involved in sensing the environmental cues by specific membrane receptors, activation of the signal transduction cascade and finally in triggering gene expression changes. These chemical reaction pathways are inherently stochastic in nature [1, 2, 3]. Stochasticity is an integral part of life at the cellular level. Processes such as gene expression,the cell cycle and signal transduction are inherently stochastic making it difficult to predict their nature. Inherent stochasticity in the external and internal environment makes cell response heterogeneous. This heterogeneous response severely limits the accuracy of estimating the input by the population of cells. However, this phenotypic heterogenity can sometimes be useful. For example, *E. coli, Staphylococcus sp*. are able to resist antibiotics due to phenotypic heterogeneity[1]. But in majority of cases, to process the cellular information correctly the cells need to be able to respond to these noisy signals appropriately implying that cells decode the environment in a probabilistic fashion.[2]. Stochasticity is often referred to as noise. This noise can either be intrinsic, i.e. a result of the process itself or, extrinsic, i.e. caused due to other processes or environmental factors[3].

Information theoretic approaches have proven to be a great asset in estimating precision of signal decoding by cellular networks[4]. The change in the output (e.g. stimulated expression of a gene) carries information about the input variable. In general, the precision with which the input value can be estimated from measuring the output improves with larger changes (i.e. dynamic range) of the output and with lower output noise. According to Cramėr–Rao inequality, the error in estimating the input from the output is bounded by the Fisher information, defined as the relative entropy change of the output distribution for an infinitesimal change in the input around a given input value[5]. Furthermore, information transmission capacity of signaling systems can also be estimated by calculating the mutual information, which is increasingly being used to characterize biochemical signaling networks[6, 7]. The mutual information measures the mutual interdependence between the input and the output distributions by calculating the relative entropy of the output distributions conditioned on the input with respect to the unconditioned output distributions[8]. In both cases, more information can be extracted about the input from the output distribution if the relative entropy change is large. The change in input is reflected in the output via the signaling pathway. To quantify the information transmission capacity we can use mutual information and/or Fisher information[9, 10].

Cells have evolved to employ multiple strategies for better information transmission namely negative feedback to reduce noise, negative regulation to amplify output dynamic range, dynamic measurement etc [11]. However, improvement of information transmission utilizing such strategies in turn imposes extra burden to the cell by accentuating energy consumption/dissipation[12]. Since, the associated chemical processes in signaling operate out of thermal equilibrium, the energy dissipation is inevitable. The close connection between information and heat dissipation in non-equilibrium processes is established on a strong theoretical footing owing to the resolution of the long standing puzzle of Maxwell’s demon through the Landauer erasure principle in the last century[13]. Entropy production rate is the measure of heat dissipation in non-equilibrium processes and entropy production rate also represents the energy consumption in many systems at steady state. In the signaling cascade operating through phosphorylation-dephosphorylation cycles, the energy consumption is essentially achieved through constant ATP consummation flux in the phosphorylation reaction and subsequent dissipation in the dephosphorylation reaction[14]. This cycle of ATP flux maintains a non-equilibrium steady state under the action of an external stimulus. Since accuracy of the response is crucial to the cell’s survival, cells have adapted to respond so that they have to consume as little energy as possible to accurately respond to information, make changes to the internal processes after learning about the external processes and do this robustly.[15]. But, how cells navigate the parameter space in designing the signaling network to achieve this trade-off of minimizing energy consumption and at the same time maintain a required accuracy level is not fully understood yet.

Here, to explore the issue, we constructed an ODE based model of the phosphorylation cascade in yeast *S. cerevisiae* pheromone response pathway incorporating random extrinsic noise in the protein levels resulting in heterogeneous cell to cell variability in response. This pathway mediates communication and ultimately mating between two haploid mating types of *S. cerevisiae*[16]. *S. cerevisiae* has two type of mating cells, MATa and MAT*α*.The *α* cells secretes the *α* pheromone which binds the G protein-coupled receptor (Ste2) and stimulates the canonical mitogen-activated protein kinase (MAPK) cascade, leading to activation of the MAPK Fus3[17, 18, 19]. Fus3, the major MAPK of the mating pathway, induces cell-cycle arrest, mating-specific changes in cell morphology by expression of mating genes including genes that encode components or regulators of the MAPK cascade[20]. Previous studies demonstrated that the pheromone pathway transmits information with high precision, as exemplified by a linear relationship between receptor occupancy and downstream responses [21] (“dose–response alignment”) and by a uniform morphological transition of cell population into a mating-competent state (“shmooing”) at a critical pheromone concentration[22, 12, 23]. Such uniformity in the output implies existence of, likely multiple, mechanisms to improve precision of information transmission within the pathway like noise reduction [24]. The pathway also consumes energy during phoshorylation of Fus3 under pheromone induction and the energy consumption is likely to increase as the negative regulator like phosphatase Msg5 is produced at higher level[14]. On the other hand, higher basal pathway activity would increase energy consumption but reduce accuracy. Thus, how these different parameters in the pathway are tuned to optimize the energy-accuracy trade-off is central to the evolutionary design of the signaling system. We show here the relation between energy consumption and accuracy of information transmission for different scenarios and compare it with experimentally fitted parameter values for the pheromone response pathway. Our study offers a theoretical framework to understand the reaction rates on the basis of evolutionary optimization.

## Results

### The reaction rates display optimality through accuracy-energy trade-off instead of only accuracy

We first consider a simple model where the Fus3 protein can make transition between inactive and active states. The activation occurs in presence of the pheromone (*α*) at a rate *α*_1_ along with a basal activation rate *α*_0_. The active state is deactivated at a rate *β*. Previous studies utilized linear response around non-equilibrium steady state to arrive at accuracy-energy trade-off analytical equation starting from fundamental principles of non-equilibrium thermodynamics. An analytical calculation provides the accuracy in estimating the input (Methods for details),

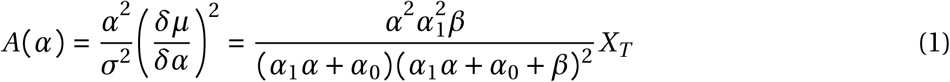

The equation 1 above shows that while we keep *α*_0_ and *α*_1_ fixed, the accuracy is zero when *β* → 0 as well as when *β* → ∞. Thus, accuracy must have a maxima at a certain value of *β*. Similarly, we also observe similar conclusion for *α*_1_ while keeping other two parameters constant (**Figure 1A**). However, we observed that the maximum value of accuracy is increasing as value of *β* increases (**Figure 1B**). In fact, if both *α*_1_ and *β* are varied simultaneously while keeping the ration 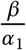 constant, the equation 1 shows that the accuracy would keep on increasing as *α*_1_ → ∞ implying that the high accuracy would be attained at high values of *α*_1_ and *β* (**Figure S1A**). Thus, the reaction rates would not reach optimum values. In case of the third parameter *α*_0_, we can clearly notice from equation 1 that the accuracy would always reduce as *α*_0_ increases. From this analysis, we can conclude that accuracy/information alone is not an ideal performance characteristic which would enable evolution to select the reaction rates optimally. Since the reaction also consumes energy in form of ATP, we performed the calculation further by incorporating the power consumption. The average power consumption by the pathway at steady state is,

**Figure 1:**
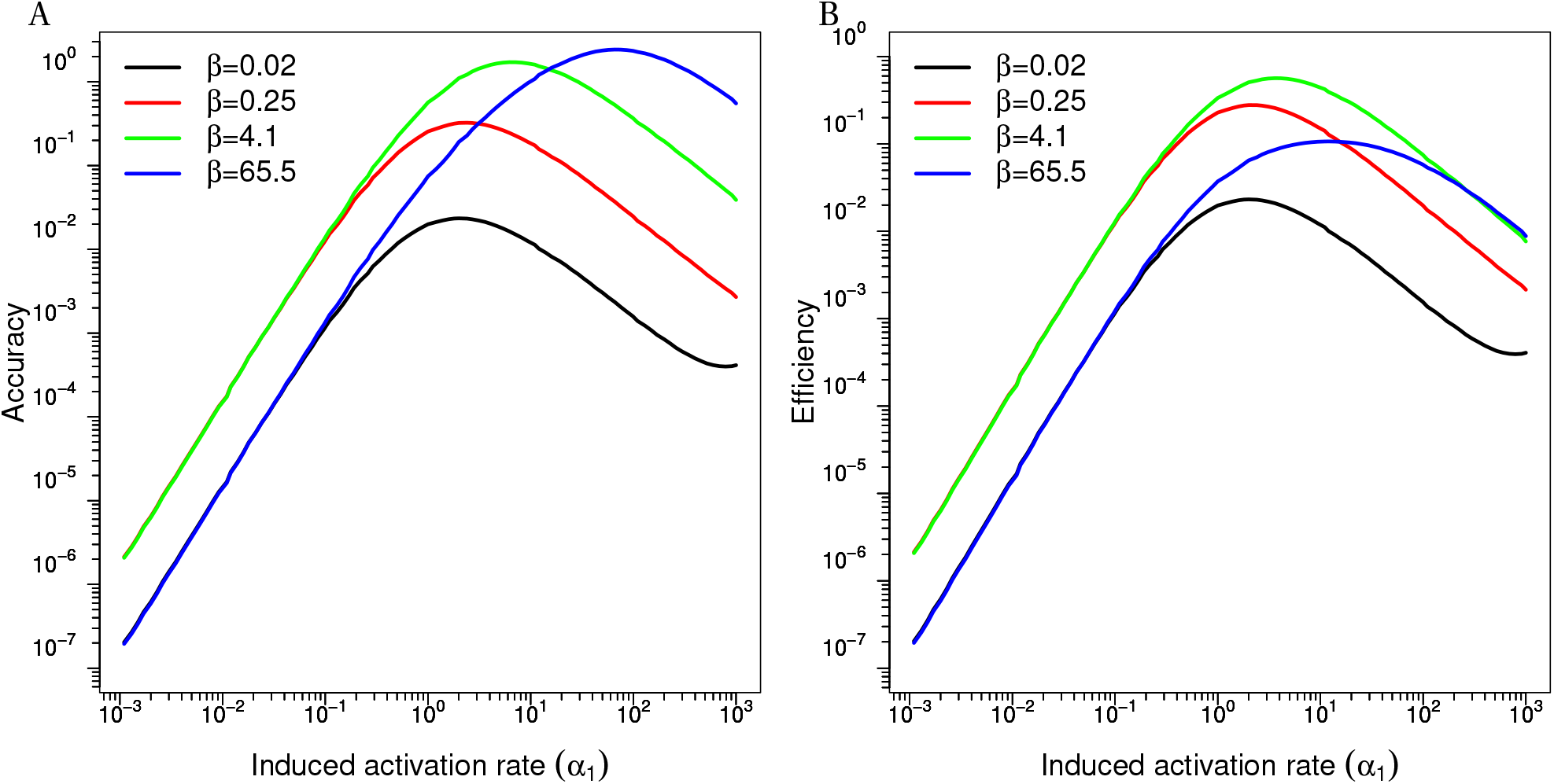
The optimization of information transmission from the analytical calculation. **(A)** The accuracy as a function the induced activation rate (*α*_1_) at different values of the deactivation rate (*β*) as indicated in the legend. **(B)** The efficiency as a function the induced activation rate (*α*_1_) at different values of the deactivation rate (*β*) as indicated in the legend.

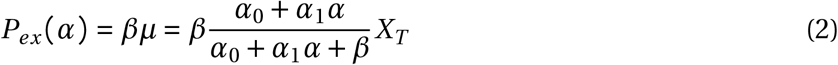

And the efficiency of performance of the signaling system is given by, (Methods for details)

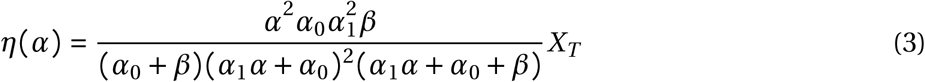

The equation 3 above for efficiency also shows optimal for *α*_1_ and *β*_1_. In fact, we can also see that by keeping 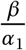 fixed, the efficiency tend to zero as *α*_1_ → ∞ which imply there exit finite values of *α*_1_ and *β* when both *α*_1_ and *β* vary simultaneously. In contrast to previous case of accuracy, now even though the maximum efficiency initially increases with *α*_1_, it starts to reduce at high value of *β* (**Figure 1B**) giving rise to an optimum set of values for both *β* and *α*_1_ (**Figure S1B**). In addition, the basal activation rate *α*_0_ also displays the optimal behaviour according to the above equation. The analysis elicits the importance of accuracy energy trade-off for evolution to navigate the parameter space and select optimal reaction rates.

### Data fitting algorithm provides estimation of the parameter values of the mathematical model

In order to explore how different parameter values like activation/deactivation rates, basal pathway activity influences the information transmission and energy consumption through and further to investigate the hypothesis presented in the previous section for a real system, we constructed a simple ODE based mathematical model. In this model, a single layer MAPK cascade of the pheromone response pathway in *S. cerevisiae* is considered. The terminal MAPK Fus3 phosphorylation is activated at a rate *k*_1_ (equivalent to *α*_1_ in the analytical model) in presence of the pheromone signal *s* as well as a basal signal *S*_*basal*_ (equivalent to *α*_0_ in the analytical model). Then the activated Fus3 is dephosphoryated at a rate *k*_2_ by a phosphatase. One component of the phosptases consists of Msg5 protein which is also transcriptionally activated by the pathway output comprising a negative feedback. The other component mediates a basal deactivation of phosphorylated Fus3 depicted as *P*_*basal*_ (equivalent to *β* in the analytical model). The phosphorylated Fus3 finally activates the transcription of a set of mating specific genes and FUS1 is one of the genes which is used as a reporter for the pathway activity. To estimate the parameter values of the simple model, we utilized a previously published experimental data set[12]. In this experiment, the pathway activity at the single cell level was measured through a GFP tagged FUS1 promoter using time-lapse fluorescence microscopy. The data set provides the dynamics of averages and variances over a population of cells at 11 different pheromone concentrations. The experiments were performed for both a WT as well as an MSG5 knock-out strains to ascertain the effect of the Msg5 mediated dephosphorylation leading to the negative feedback loop discussed earlier. We first implemented the fitting algorithm on the average dose response curves at first four different time points and all the 11 pheromone concentrations for the MSG5 knock out strain (**Figure 2A**). This provides us with the average of all the parameter values excluding the Msg5 specific parameter values. Finally, the WT dose response curves are fitted keeping the previously obtained parameter values fixed and only varying the parameter values associated with Msg5 mediated deactivation (**Figure 2A**). Through this method, we were able to fit the dose-response curves of WT and MSG5 knock-out strains simultaneously (estimated parameter values are provided in the appendix). Since, the pathway activity is measured indirectly through the FUS1 gene expression output, we conducted a sensitivity analysis to validate whether the upstream parameter values have significant contribution to the curve fitting. Although the downstream parameters like the GFP production and degradation rates expectedly have higher sensitivities, the key parameters of the core pathway e.g. *k*_1_, *S*_*basal*_ and *P*_*basal*_ also display significant sensitivities (**Figure 2B**). Next, we would explore how the information transmission through the pathway is optimized by varying these key three parameters around their estimated values.

**Figure 2:**
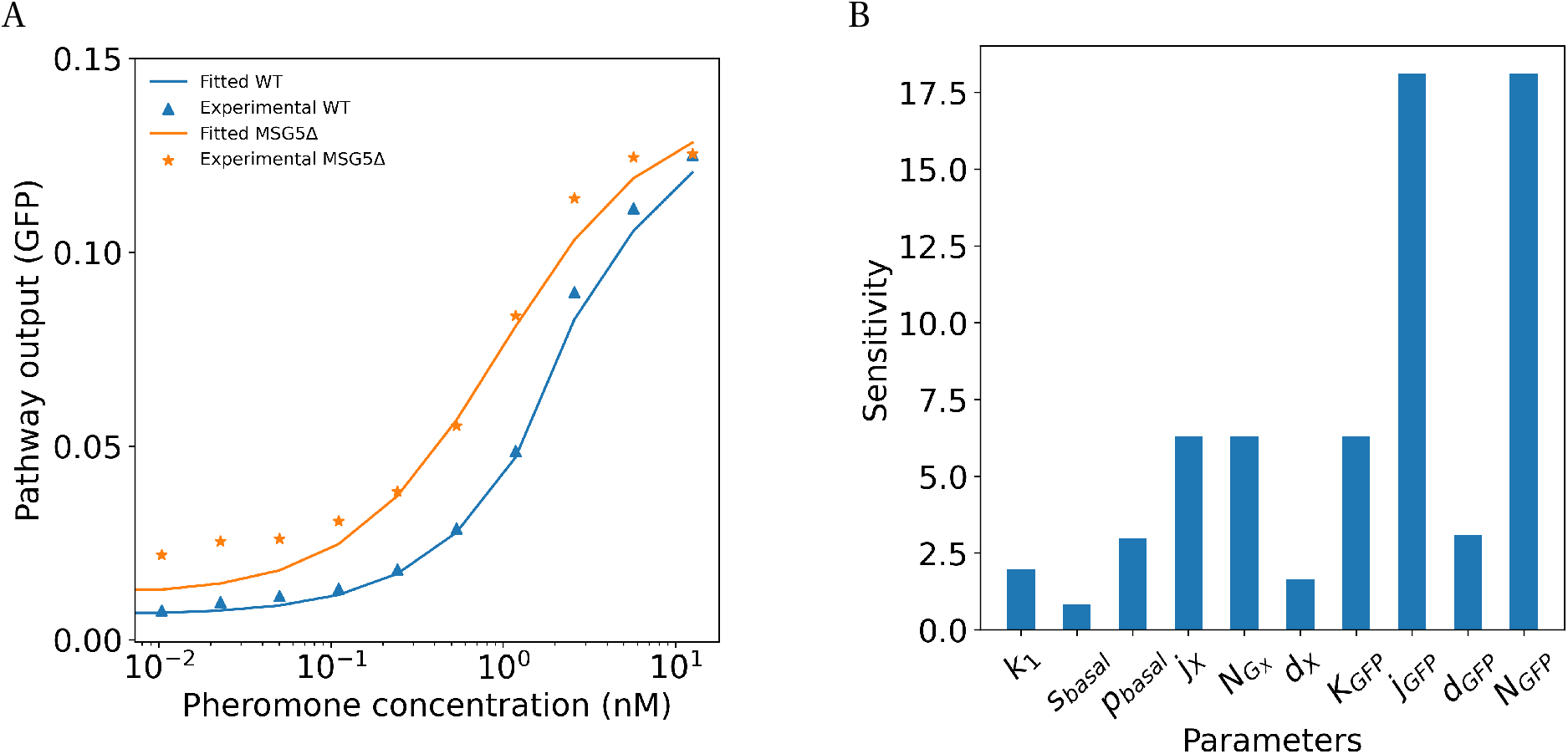
Reaction rates are estimated by fitting the experimentally measured dynamics of the pathway output with the mathematical model. **(A)** The fitted curves along with the experimentally measured GFP output values at the third time index are shown for the WT and the MSG5Δ strain as indicated. **(B)** Sensitivity analysis of all the parameters for the MSG5Δ variant.

### Higher basal dephophorylation rate reduces the basal pathway activity and forces response to be induced at higher pheromone doses

First, in order to investigate the effect of basal dephophorylation rate on the responses, we generate dose response curves at different pheromone concentration by performing simulation by varying *P*_*basal*_ around the estimated value (PE) while keeping the Msg5 concentration at zero, resembling the MSG5 knock-out strain (**Figure 4A**). Initially, the basal pathway activity is observed to be gradually reduced as the *P*_*basal*_ increases while the saturation levels of Fus3 activity do not change significantly resulting in amplification of overall dynamic ranges. However, a very high value of *P*_*basal*_ reduces the saturation level ultimately leading to compression of the dynamic range.

**Figure 3:**
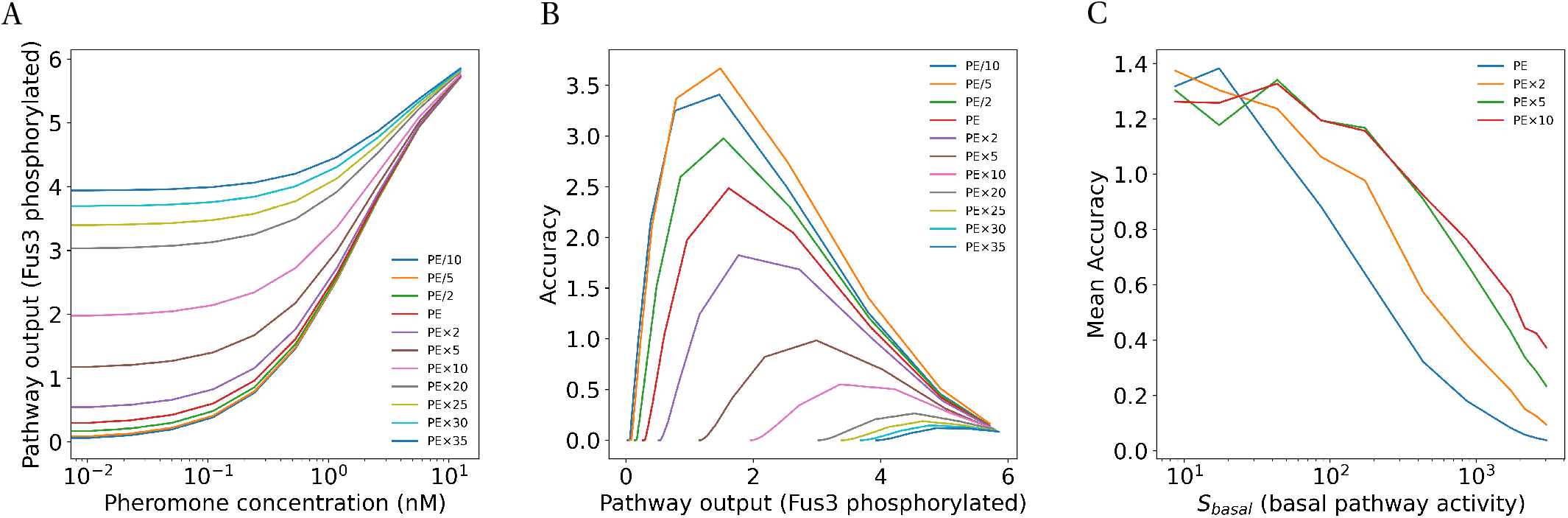
The accuracy degenerates with higher basal pathway activity (*S*_*basal*_). **(A)** Dose-Response Curves for different values of *S*_*basal*_ where PE corresponds to the estimated parameter value of *S*_*basal*_ while keeping other parameter values fixed at estimated values. **(B)** The simulation results for accuracy vs pathway output at the Fus3 phosphorylation level for different values of *S*_*basal*_ where PE corresponds to the estimated parameter value of *S*_*basal*_ while keeping other parameter values fixed at estimated values.**(C)** Mean accuracy vs *S*_*basal*_ at four different values of the induced activation rate (*k*_1_) as indicated in the legend while keeping other parameter values fixed at estimated values. PE corresponds to the estimated parameter value of *k*_1_.

**Figure 4:**
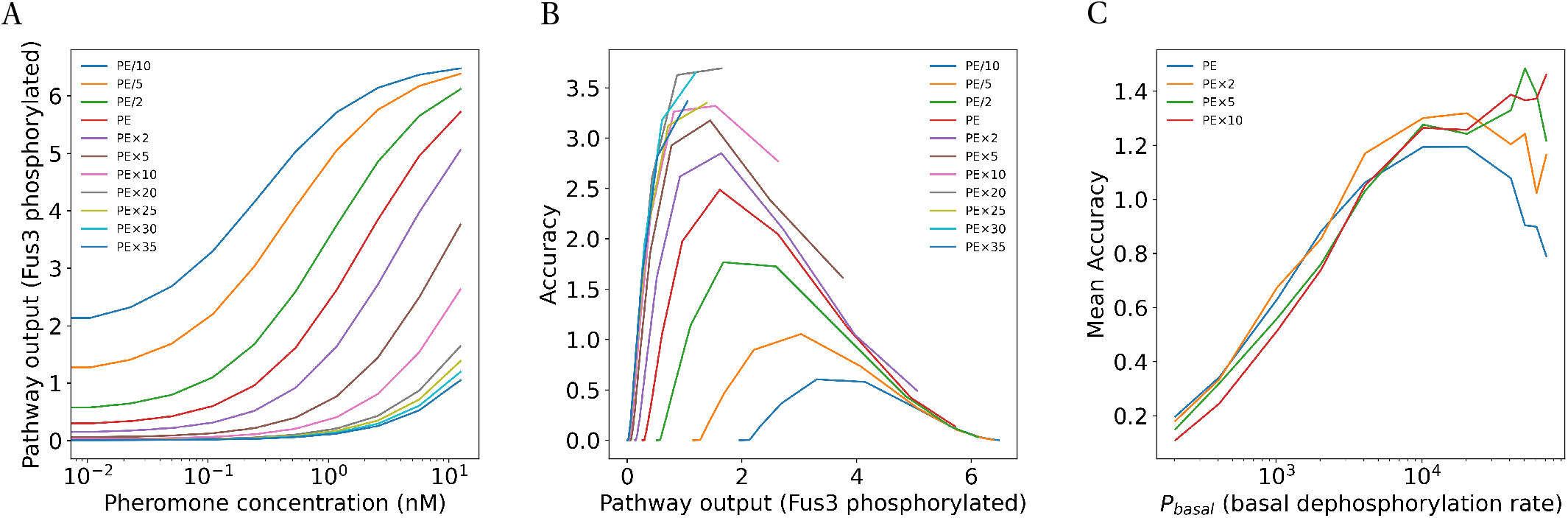
The accuracy displays an optimum for the basal deactivation rate (*P*_*basal*_). **(A)** Dose-Response Curves for different values of *P*_*basal*_ where PE corresponds to the estimated parameter value of *P*_*basal*_ while keeping other parameter values fixed at estimated values. **(B)** The simulation results for accuracy vs pathway output at the Fus3 phosphorylation level for different values of *P*_*basal*_ where PE corresponds to the estimated parameter value of *P*_*basal*_ while keeping other parameter values fixed at estimated values.**(C)** Mean accuracy vs *P*_*basal*_ at four different values of the induced activation rate (*k*_1_) as indicated in the legend while keeping other parameter values fixed at estimated values. PE corresponds to the estimated parameter value of *k*_1_.

### Accuracy and mutual information show an optimum with respect to *P_basal_*

Next, we introduced noise through adding cell to cell variability in the protein expression (Methods). The intrinsic fluctuation due to stochasticity is neglected since the intrinsic noise has been shown to be low for this pathway [25]. In presence of noise, the information transmitted through the pathway can be calculated by quantifying either the accuracy of estimating the input from the output or the mutual information between the input and output. The accuracy is bounded by the fisher information according to Cramėr–Rao inequality. We calculated the accuracy at different pheromone concentration by varying *P*_*basal*_ values(**Figure 4B**). We observed that accuracy is very small at low doses and gradually increase. But, it becomes small again at saturating dose as the response does not change even if the dose increases. The maximum accuracy is achieved at intermediate pheromone concentrations. To quantify the overall accuracy at a particular dose, the average accuracy is calculated. It is noticed that the average accuracy as well as mutual information improve as the *P*_*basal*_ increases owing to the enhanced dynamic range but it starts to drop at high value of *P*_*basal*_ due to reduced saturation level (**Figure 4C**;**Figure S2B**). Thus, when other parameter values are fixed, the *P*_*basal*_ exhibits an optimum where the accuracy is maximum.

### Higher basal pathway activity impairs accuracy

Next, we explored the effect of basal pathway activity by varying *S*_*basal*_. Now, simulations were performed by changing values of *S*_*basal*_ while keeping other parameter fixed. Clearly, if the value of *S*_*basal*_ increases, the basal pathway activity elevates while the saturation level remains the same leading to a compression in the dynamic range. (**Figure 3A**). The compression in the dynamic range gives rise to low accuracy and mutual information (**Figure 3B**). Thus, the average accuracy and mutual information gradually diminishes with higher *S*_*basal*_ (**Figure 3C**;**Figure S2A**).

### Higher activation rate improves accuracy but accuracy degenerates at high activation rates

Finally, to explore the effect of the pheromone induced activation rate, we varied the value of *k*_1_ and simulate the pathway output. In this case, initially at a very low *k*_1_ values, the pathway is not induced at all, only operating at basal level (**Figure 5A**). The mean accuracy is subsequently low (**Figure 5B**). As the value of *k*_1_ is increased, the saturation level starts to grow, effectively amplifying the dynamic range of the dose response curve. Thus, the overall accuracy also improves(**Figure 5C**). However, at very high activation rates, the pathway gets activated even at very low doses giving rise to loss of accuracy and information (**Figure 5C**;**Figure S2C**). Similar to the basal dephosphorylation rate, the activation also exhibits an optimum with respect to the accuracy. Thus, we observed that while other parameters remain fixed, both the *P*_*basal*_ and *k*_1_ exhibit optimum values where the accuracy is maximum. However, as observed before, the maximum accuracy increases as the value of *P*_*basal*_ increases. The basal activity mediated by *S*_*basal*_ on the other hand does not show any such optimum. From an evolutionary point of view, if achieving high information transmission capacity is desired, high parameter values will be selected for *P*_*basal*_ and *k*_1_ whereas very low value of *S*_*basal*_ would be desirable. But, according to the experimentally obtained estimation, the estimated value of *P*_*basal*_ and *k*_1_ are quite low. The estimated value of *S*_*basal*_ is quite high. Furthermore, all the parameters would change simultaneously during evolution in contrast to the scenario considered here where all parameters were kept fixed to calculate the optimum value of either *k*_1_ or *P*_*basal*_.

**Figure 5:**
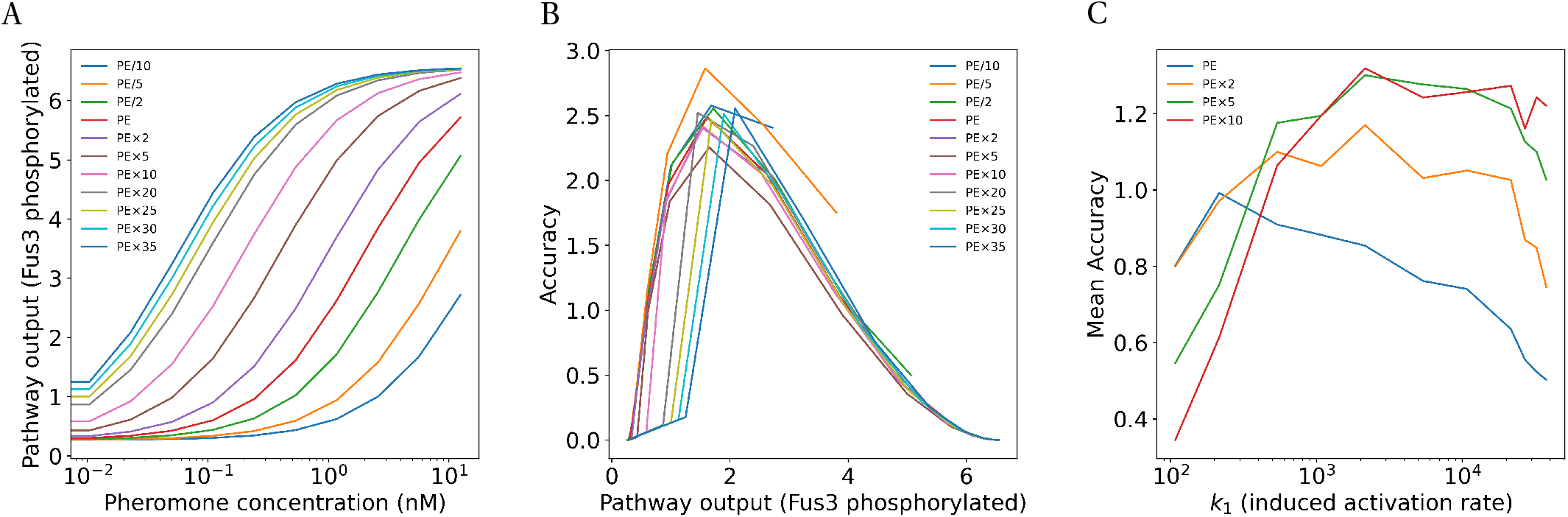
The accuracy display an optimum with respect to the induced activation rate (*k*_1_). **(A)** Dose-Response Curves for different values of *k*_1_ where PE corresponds to the estimated parameter value of *k*_1_ while keeping other parameter values fixed at estimated values. **(B)** The simulation results for accuracy vs pathway output at the Fus3 phosphorylation level for different values of *k*_1_ where PE corresponds to the estimated parameter value of *k*_1_ while keeping other parameter values fixed at estimated values.**(C)** Mean accuracy vs *k*_1_ at four different values of the basal deactivation rate (*P*_*basal*_) as indicated in the legend while keeping other parameter values fixed at estimated values. PE corresponds to the estimated parameter value of *P*_*basal*_.

### The optimality is not observed in 2-dimension

As observed in the previous section that the values *P*_*basal*_ and *k*_1_ show optimum values individually, we performed simulations by varying both *P*_*basal*_ and *k*_1_ simultaneously and calculated the average accuracy and mutual information for each set of values (**Figure 6A**). In fact, the accuracy and information keep on increasing along the diagonal. In corroboration with previous observation, for a particular value of *k*_1_ along a particular column the *P*_*basal*_ has an optimum value where accuracy is maximum. Similar result can be observed along a row where *k*_1_ is fixed. But the maximum accuracy value keeps on increasing along the diagonal. Essentially, the 1-dimension curves can be thought of a projection of 2-D contour on 1-D. Thus, there seems to be no optimum values of *k*_1_ and *P*_*basal*_ where accuracy or information is maximum. The desirable values of *k*_1_ and *P*_*basal*_ can attain any high values keeping the ratio same. Similar results are observed for the mutual information (**Figure S3C**). We further performed simulations by varying *S*_*basal*_ and *P*_*basal*_ simultaneously (**Figure 6B**;**Figure S3A**). As expected, the *S*_*basal*_ and *P*_*basal*_ do not show any optimum values since we already demonstrated that *S*_*basal*_ does not posses any optimal value even in 1-dimension. A natural consequence of this analysis is the hypothesis that the accuracy or information is not the only determining factor in naturally selecting the parameter values. In that case, the evolution would navigate towards very high values of *P*_*basal*_ and *k*_1_ while keeping the value of *S*_*basal*_ as low as possible.

**Figure 6:**
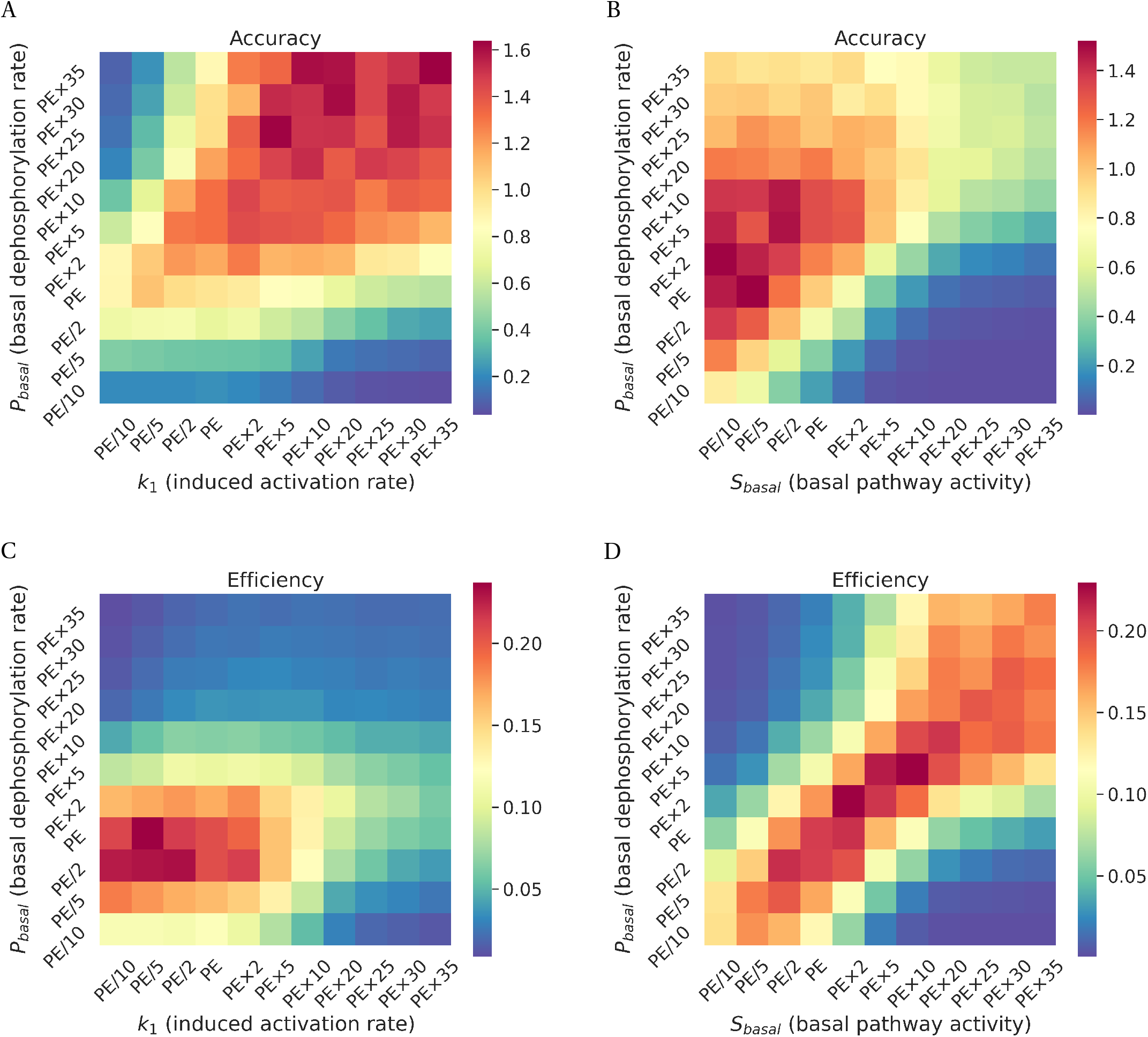
Optimality of the reaction rates with respect to the accuracy-energy trade-off. **(A)** Heatmap shows the accuracy as function of *P*_*basal*_ and *k*_1_. The PE on x-axis corresponds to estimated value of the induced activation rate (*k*_1_) while The PE on y-axis corresponds to estimated value of the basal deactivation rate (*P*_*basal*_). **(B)** Heatmap shows the accuracy as function of *S*_*basal*_ and *P*_*basal*_. The PE on x-axis corresponds to estimated value of the basal pathway activity (*S*_*basal*_) while The PE on y-axis corresponds to estimated value of the basal deactivation rate (*P*_*basal*_). **(C)** Heatmap shows the efficiency as function of *P*_*basal*_ and *k*_1_. The PE on x-axis corresponds to estimated value of the induced activation rate (*k*_1_) while The PE on y-axis corresponds to estimated value of the basal deactivation rate (*P*_*basal*_). Heatmap shows the efficiency as function of *S*_*basal*_ and *P*_*basal*_. The PE on x-axis corresponds to estimated value of the basal pathway activity (*S*_*basal*_) while The PE on y-axis corresponds to estimated value of the basal deactivation rate (*P*_*basal*_).

### A trade-off between accuracy and energetic cost can explain the choice of reaction rates

The phosphorylation of Fus3 by the upper kinase occurs by moving a phosphate group on ATP to one of the serine/theonine residues of Fus3, converting an ATP molecule to ADP. The cells energy budget effectively drains out due to this continuous cycle. The rate of energy consumption would increase with higher activation/deactivation rates. Intuitively, we can now imagine that even though the accuracy improves by making the activation/deactivation rate higher, the pathway will consume more power posing a fundamental trade-off for the cell. In order to explore this trade-off, we defined an efficiency for the pathway which naturally appears in equation for power-accuracy trade-off by a linear response theory calculation (Methods).

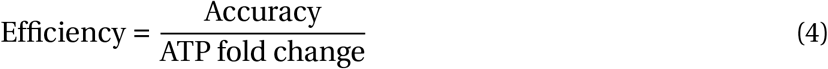

Where ATP fold change is the fold increase in ATP consumption rate from the basal ATP consumption rate. This formulation essentially makes efficiency a unit-less number. From the simulation, we now calculated the value of efficiency at each dose and finally took an average over the doses. We observe that the *P*_*basal*_ and *k*_1_ values shows an optimum low values close to the estimated value where efficiency is maximum (**Figure 6C**). The *S*_*basal*_ also exhibits a finite value close to the estimated value (**Figure 6D**). The analysis suggests that the pheromone response pathway is not optimized to maximize information transmission but it needs to balance the energetic cost with information transmission for sustainability. It highlights the importance of both information transmission and energetic cost in determining the reaction rates of the signaling pathway. The analysis till now was performed for the MSG5 knock-out strain to arrive at the conclusion. We repeat the same analysis by adding the feedback loop in the pathway. Considering only accuracy or information, the inability to obtain an optimum value when *P*_*basal*_ and *k*_1_ vary simultaneously appears here as well (**Figure 7A**;**Figure S3D**). The *S*_*basal*_ does not show any optimum as expected (**Figure 7B**;**Figure S3B**). Finally the optimum values of *k*_1_, *P*_*basal*_ and *S*_*basal*_ arrive close to the estimated value when we consider the trade-off between accuracy and energetic cost (**Figure 7C, 7D**).

**Figure 7:**
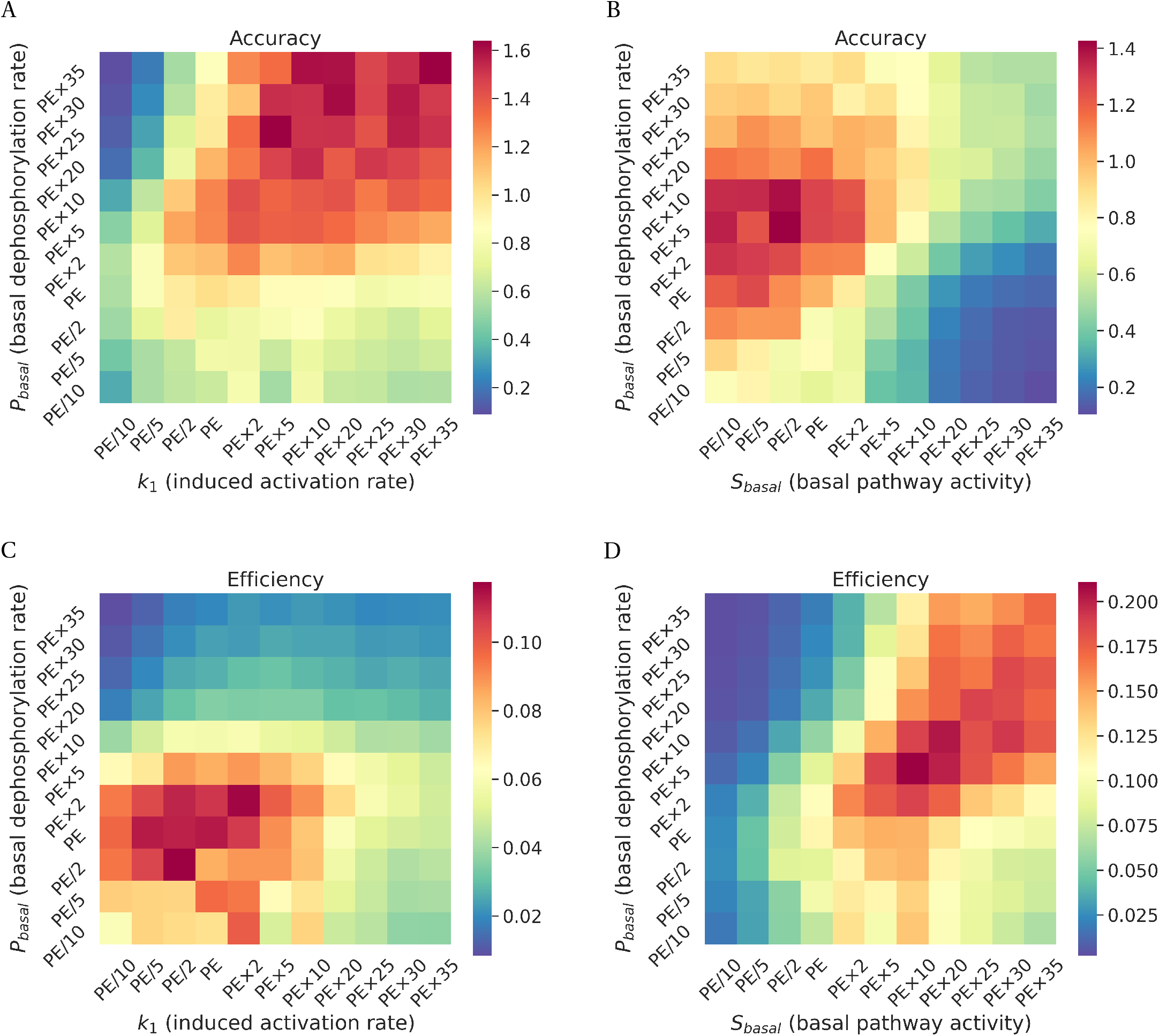
Optimality of the reaction rates with respect to the accuracy-energy trade-off by including the negative feedback. **(A)** Heatmap shows the accuracy as function of *P*_*basal*_ and *k*_1_. The PE on x-axis corresponds to estimated value of the induced activation rate (*k*_1_) while The PE on y-axis corresponds to estimated value of the basal deactivation rate (*P*_*basal*_). **(B)** Heatmap shows the accuracy as function of *S*_*basal*_ and *P*_*basal*_. The PE on x-axis corresponds to estimated value of the basal pathway activity (*S*_*basal*_) while The PE on y-axis corresponds to estimated value of the basal deactivation rate (*P*_*basal*_). **(C)** Heatmap shows the efficiency as function of *P*_*basal*_ and *k*_1_. The PE on x-axis corresponds to estimated value of the induced activation rate (*k*_1_) while The PE on y-axis corresponds to estimated value of the basal deactivation rate (*P*_*basal*_). **(D)** Heatmap shows the efficiency as function of *S*_*basal*_ and *P*_*basal*_. The PE on x-axis corresponds to estimated value of the basal pathway activity (*S*_*basal*_) while The PE on y-axis corresponds to estimated value of the basal deactivation rate (*P*_*basal*_).

## Discussion

Our study provides a theoretical formalism to understand the evolutionary choice of the reaction rates of a signaling pathway. The analysis is essentially based on the fundamental principle of information-thermodynamics connection[13]. We show that the information transmission cannot be be only determining factor for evolution to act on. The energetic cost in the transmitting signal also plays a crucial role in evolution. Here, we specifically analysed the trade-off between the accuracy and energetic cost around the estimated values of the pathway. In fact, the optimum values arrive close to the estimated value only when the energetic cost is also incorporated in the picture in addition to the information transmission. Consideration of only information transmission does not fully explain low values of activation/deactivation rates. Although, the fraction of energy investment in the signaling cycle is small, previous experiment has demonstrated that this small amount of cost can have significant consequences for growth of single cell organisms in evolutionary time scales[12, 26]. The role of energetic cost in cellular signaling systems has been implicated in previous studies [27, 28, 26, 12]. Apart from cellular signaling, the similar consideration in shaping the rates in reaction networks was investigated for error proof reading in translation [29], as well as in copying biochemical systems [30].

In this study, we primarily used accuracy to determine the performance of the signaling system. The accuracy is defined as the fisher information at a particular input [31, 12]. The conventionally used mutual information [4, 9] shows similar results. The choice of accuracy is contingent on the fact that the accuracy can be quantified at a particular input, where as the mutual information can only be defined for a set of inputs. Since, energy consumption changes with the input, the individual efficiency defined here can only be calculated at a particular input and then taking average over all the inputs. Additionally, the fisher information naturally arises in connecting the power consumption to the accuracy using linear response theory discussed here [32, 33, 34]. In fact, a strong connection between mutual and fisher information has already already been established mathematically [5].

For this particular signaling pathway the estimation of the pheromone concentration is crucial for mating decision which make accuracy a natural choice for evolution to act on. However, we need to be careful applying the principle in general cases. It may happen that the primary aim of the signaling system may not be estimating the concentration of the input, but rather estimating the category of the input. Thus, the determining fitness factor must be chosen with care. Finally, the analysis should be extensively performed in multiple dimension. As we have illustrated the parameter values may display optimum in 1 dimension but not 2 dimensions. Similarly, optimum obtained in 2 dimensions may not be observed in higher dimension.

Qualitatively, we are able to demonstrate the importance of the energy accuracy trade-off. In spite of the fact the simulation is able to capture the estimated reaction rates close to the optimum value, they are not exactly the same. The optimum value also depends on the range of the pheromone concentrations on which the analysis is performed. It is possible that we exactly don’t know the range of the pheromone concentrations the cells encounter in the natural environment and the pathway reaction rates would cater to the natural environment. Additionally, the parameter values are estimated through the gene expression output rather then directly at the phosphorylation level output which will, to some extent, obscure parameter estimation at the upstream level.

## Methods

### Analytical expression for accuracy and efficiency

For a non-equilibrium stochastic process of a random variable x, the steady state probability distribution can be denoted by *ρ*(*x, α*) where *α* is the control parameter. Following Hatano-Sasa formulation [33] for transition between non-equilibrium steady states, the non-equilibrium free energy can written as

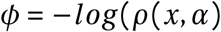

The conjugate force corresponding to the control parameter *α* is given by

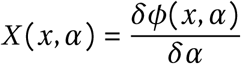

For a small change in the value of *α* to *α* + *δα*, the average change in the conjugate force around the steady state value can be derived using linear response theory, 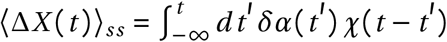

Where,

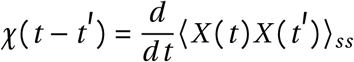

The thermodynamic internal energy due to the process can be represented as

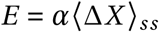

and the corresponding excess power generated due to the process is [32]

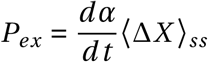

By integrating by parts, we can finally get

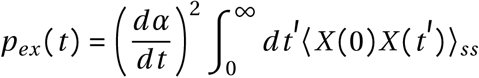

For the single stage phosphorylation-dephosphorylation reaction system considered here, the steady state equation for phosphorylated protein concentration is given by,

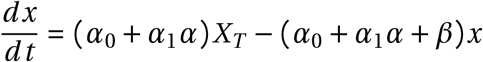

The Langevin equation at steady state is,

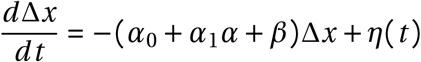

The steady state distribution around the steady state is given by,

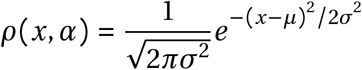

Following the steps above, it can be shown that,

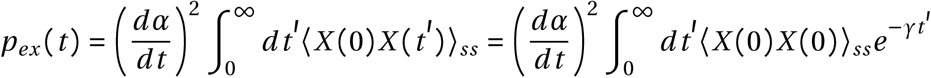

Since, the fisher information *F* at steady state is defined as,

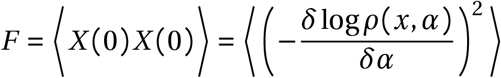

Thus, finally the calculation leads to a power-accuracy trade-off [35].

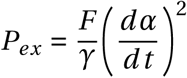

According to Cramėr–Rao inequality 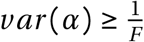 which imply that,

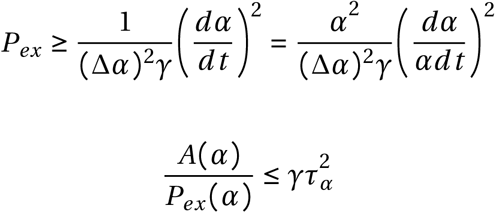

Here, 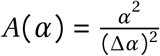 is the accuracy of estimating the signal and inverse of 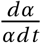 represents time scale of the signal change *τ*_*α*_. In order to make the efficiency unitless, we multiply the equation by *p*_*ex*_ (0), i.e. power consumtion when *α* = 0. The efficiency *η*(*α*) is defined as

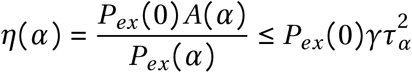

The efficiency is the accuracy obtained in estimating the signal *α* per unit of fold change in the power consumed 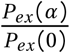. For the case of phophorylation reaction,

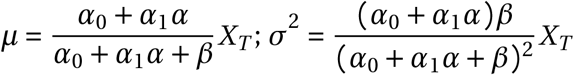

The accuracy [28, 12] is the fisher information with respect to the input *α* and if we assume that total number of the sensing molecule *X*_*T*_ is large, it can be shown to be given by,

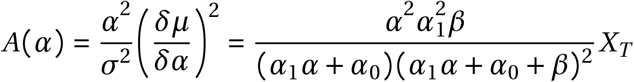

The total accuracy can be quantified by summing over the input signals, 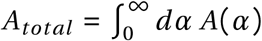. The average power consumption,

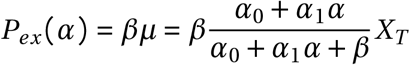

And the efficiency,

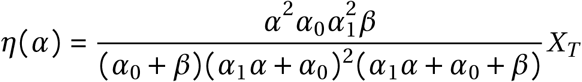

The total efficiency over the input can be quantified by 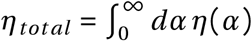

### Detailed mathematical model for simulation study

We consider one-step phosphorylation and cascade of the model pathway, where the receptor pheromones activate kinase Fus3 and which induces the activity in Msg5 and GFP. The system is modelled as a set of Ordinary Differential Equations. We build a mathematical model where *X* is phosphorylated and thus activated by pheromone *s*. This *X* after phoporylation becomes *X* ^*p*^ which is a promoter of GFP as well as *P*. *P* then activates the phosphatase which in turn promotes the dephosphorylation of *X* ^*p*^ back to *X*. This system is modelled using simple Ordinary Differential Equations (ODEs). The simulations were performed using the ode15s solver in MATLAB 2019b for a population of 500 cells. Here *X* represents inactive Fus3, *X* ^*p*^ represents acitve or phophorylated Fus3 and *P* represents Msg5 proteins respectively. Given below are the ODEs for different proteins.

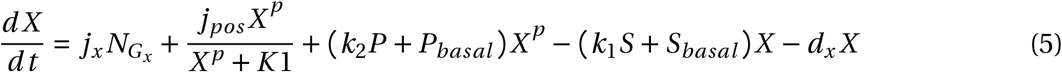

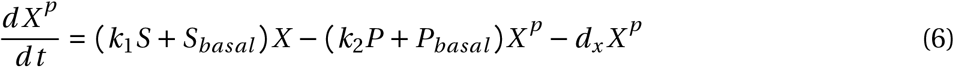

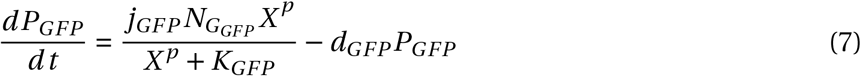

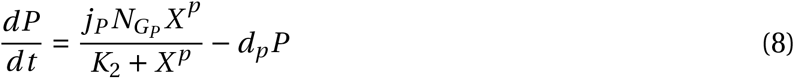

For differing values of the parameters *P*_*basal*_, *k*_1_ and *S*_*basal*_ we simulate a population of 500 cells with normal noise introduced in the production rates of all proteins. The values of aforementioned parameters were changed by making the fitted value 2-fold, 5-fold, 10-fold, 20-fold, 25-fold, 30-fold and 35-fold of itself and dividing the fitted value by 2, 5 and 10.

### Calculating mutual information and accuracy

To calculate mutual information between the input and output (in our case the input being the pheromone concentration, also known as the signal *S* and the output being the response *R*) we need to look at the probability distribution of both *S* and *R* and thier joint probability distribution. However, we only know the conditional probability, *P* (*R*|*S*). Thus using the below equation

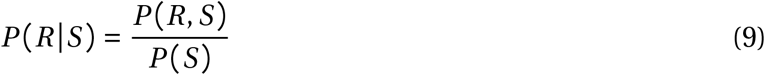

We can rewrite Equation 9 as

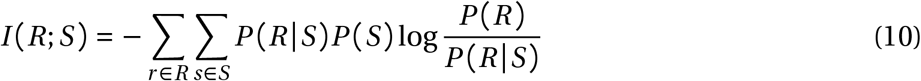

During the experiments, we vary the pheromone concentration to one of 11 values. As we run the same number of experiments for each one of the values we get a uniform distribution for the input. We then proceed to add noise in the production rate of all downstream proteins in our model and simulate the output *R* for different noise 500 times. The values of *R* are then arranged into 40 bins of a histogram. We use this histogram to estimate the probability distribution of the response *R* for each one of the 11 values of the signal *S* Thus we are able to find the mutual information by finding the entropy of the output *R* and subtracting the conditional entropy of of output *R* given input *S* using the equation stated below.

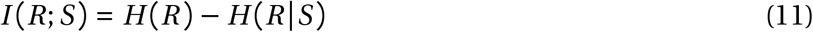

To measure the accuracy in the dose-response curve we measure the derivative at every dose of the dose-response curve and the standard deviation in the response. The accuracy is given as

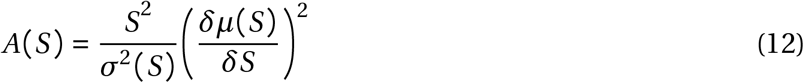

where,

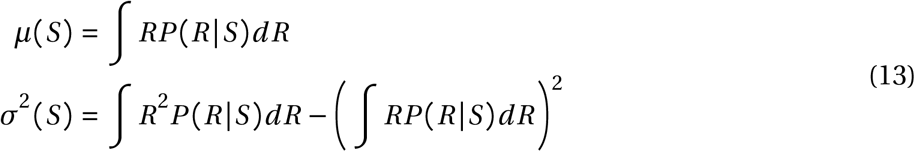

As the experimental data used is the single cell sequence data, we can obtain the *μ*(*S*) and *σ* (*S*) by using predefined functions in Python[36]. Using scipy[37] package’s B-spline interpolation, we are able to obtain simulated dose-response curves. The interpolated curves, although discrete, can be used to obatin the accuracy. The derivative of the dose response curve can be given by

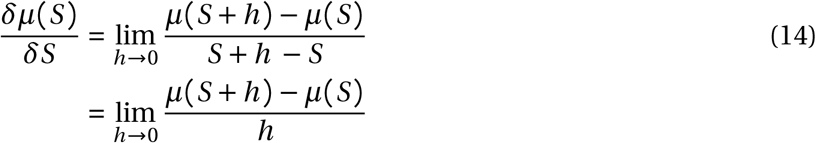

For practical purposes *h* is chosen in the order of 10^−5^ while obtaining the simulated dose-response curve. By substituting the result of Equation 14 in Equation 12 we obtain the accuracy for each dose.

### Parameter Estimation

The parameters for the mathematical model were estimated using the genetic algortihm (ga) function in MATLAB’s global optimization toolbox. The loss function was set to minimize the root mean squared error (RMSE) between the experimental dose response curve and the predicted dose response curve from the model. The parameters were first estimated for the MSG5 knock-out strain. Then keeping these parameters constant the rest of the parameters pertaining to Msg5 activity in the WT strain were estimated.

### Sensitivity Analysis

To find the senstitivity of the dose response curve to the change in the parameter values we change a particular parameter while keeping all the other parameter values the same as they were obtained from parameter estimation. The parameter in question is changed by multiplying or dividing it by 2, 5 and We then take the root mean squared error (RMSE) between the dose-response curve obtained after changing the parameter and the experimental dose-response curve let it be RMSE_change_. An average of all RMSEs for changes in the same parameter is taken, Mean RMSE_change_.

Let RMSE between the experimental dose-response curve and simulated dose-response curve for all parameter values being the same as obtained from the parameter estimation be, RMSE_*or iginal*_. Therefore we define sensitivity, or a particular parameter, as

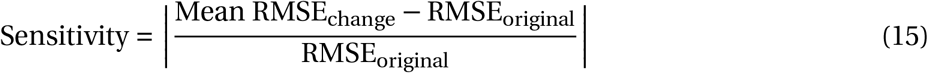

## Acknowledgements

Authors thank the Department of Biotechnology (No. BT/RLF/Re-entry/32/2017), Government of India for funding.

## Contributions

B.G. conceived the study. A.M. performed the simulation studies. A.M. and B.G. wrote the manuscript.

## Appendix

### Figures

**Figure S1:**
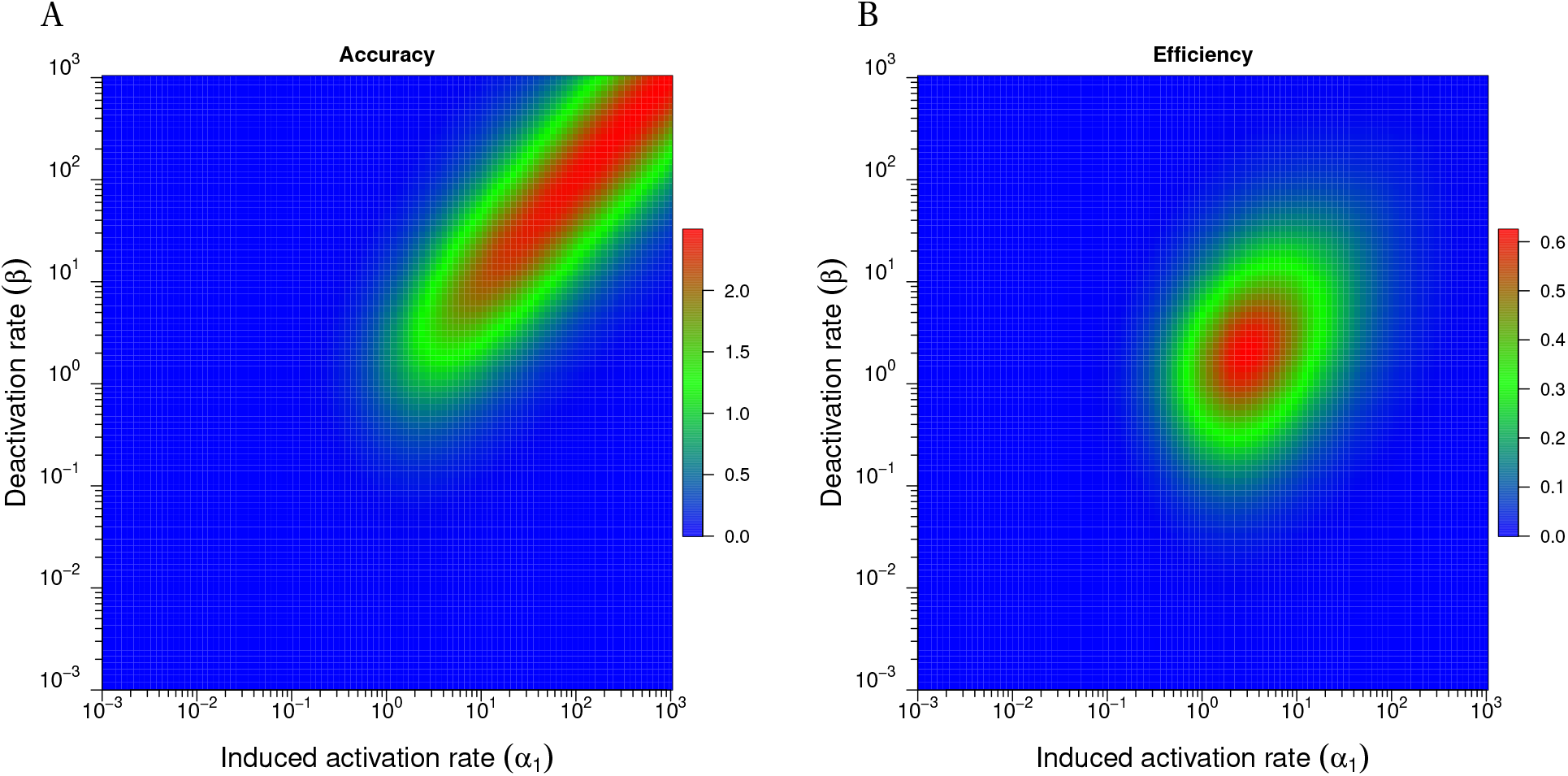
The optimization of information transmission from the analytical calculation in 2-dimension. **(A)** The heatmap displays the accuracy as function of the induced activation rate (*α*_1_) and the deactivation rate (*β*) **(B)** The heatmap displays the efficiency as function of the induced activation rate (*alpha*_1_) and deactivation rate (*beta*)

**Figure S2:**
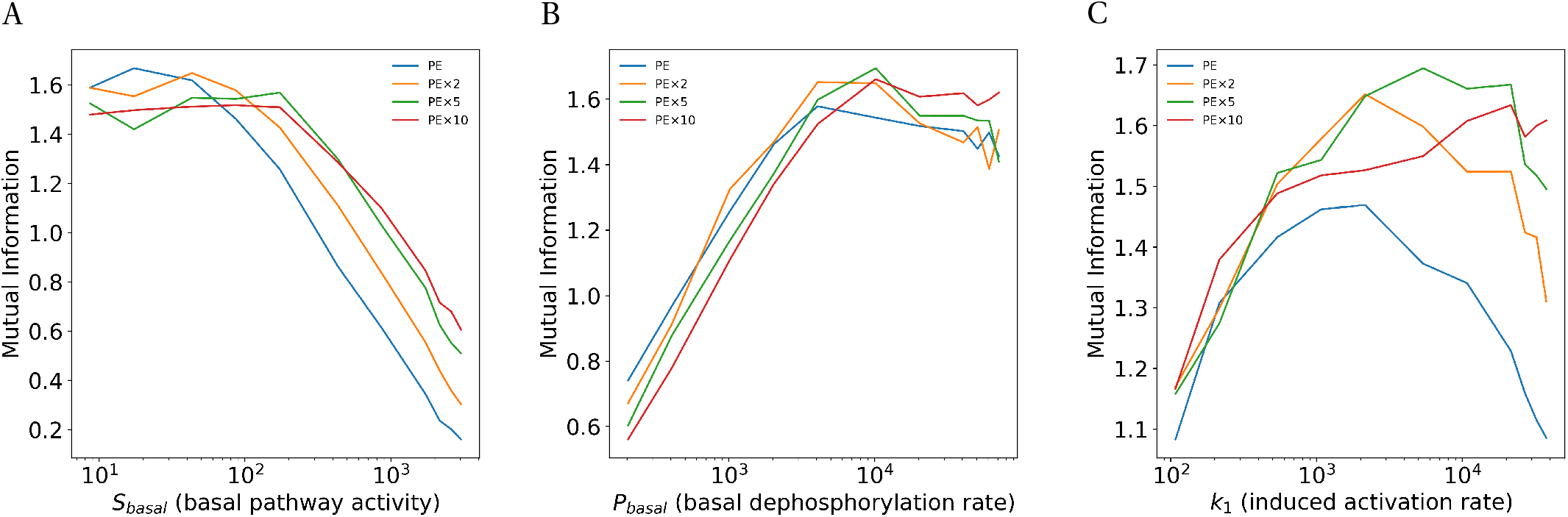
**(A)**Mutual information vs *S*_*basal*_ at four different values of the induced activation rate (*k*_1_) as indicated in the legend while keeping other parameter values fixed at estimated values. PE corresponds to the estimated parameter value of *k*_1_.**(B)**Mutual information vs *P*_*basal*_ at four different values of the induced activation rate (*k*_1_) as indicated in the legend while keeping other parameter values fixed at estimated values. PE corresponds to the estimated parameter value of *k*_1_. **(C)**Mutual information vs *k*_1_ at four different values of the induced activation rate (*P*_*basal*_) as indicated in the legend while keeping other parameter values fixed at estimated values. PE corresponds to the estimated parameter value of *P*_*basal*_.

**Figure S3:**
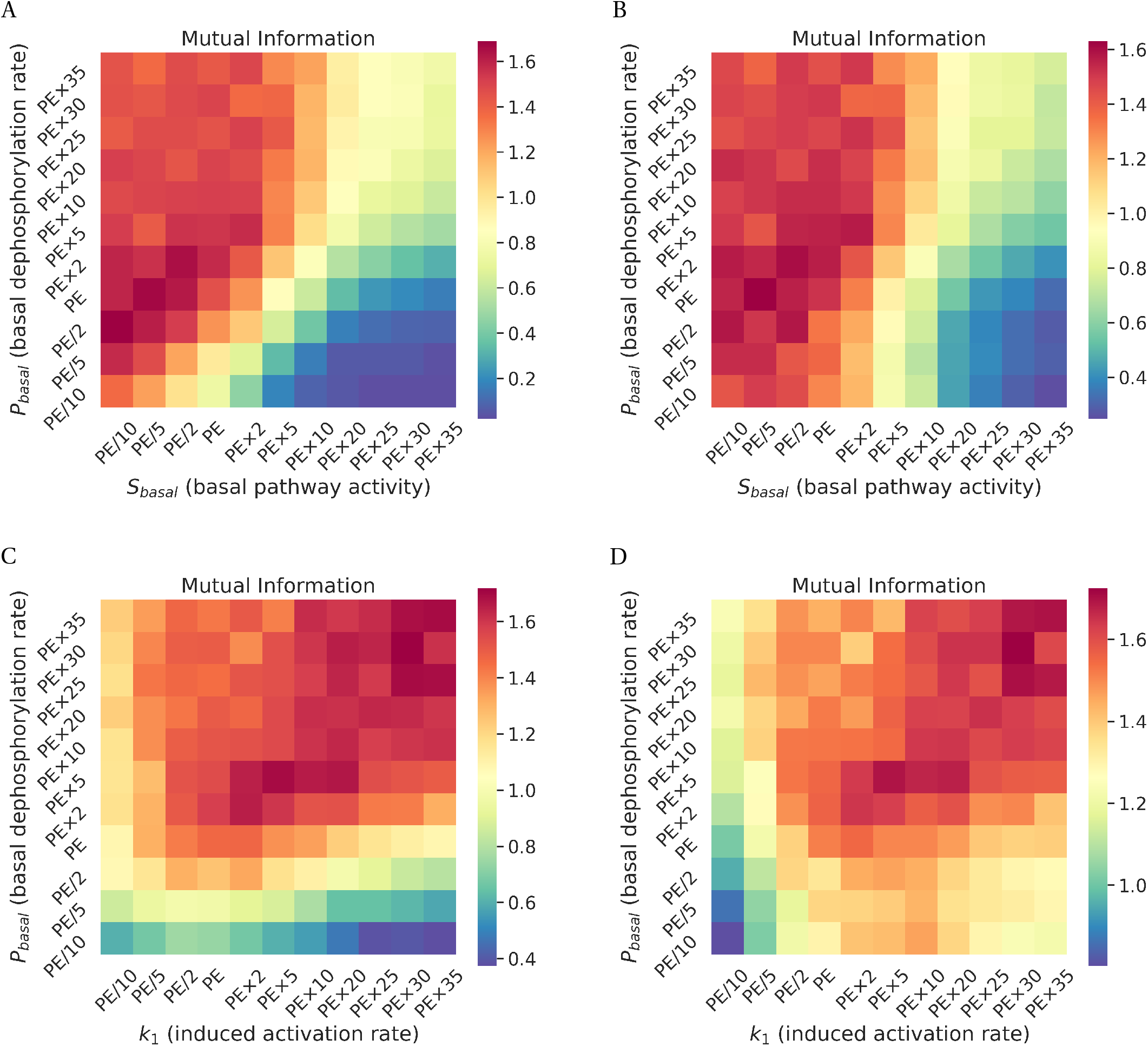
**(A)** Heatmap shows the mutual information as function of *S*_*basal*_ and *P*_*basal*_. The PE on x-axis corresponds to estimated value of the induced activation rate (*S*_*basal*_) while The PE on y-axis corresponds to estimated value of the basal deactivation rate (*P*_*basal*_). **(B)** Heatmap shows the accuracy as function of *S*_*basal*_ and *P*_*basal*_ for WT in presence of the induced feedback loop. The PE on x-axis corresponds to estimated value of the basal pathway activity (*S*_*basal*_) while The PE on y-axis corresponds to estimated value of the basal deactivation rate (*P*_*basal*_). **(C)** Heatmap shows the mutual information as function of *k*_1_ and *P*_*basal*_. The PE on x-axis corresponds to estimated value of the induced activation rate (*k*_1_) while The PE on y-axis corresponds to estimated value of the basal deactivation rate (*P*_*basal*_). **(D)** Heatmap shows the accuracy as function of *k*_1_ and *P*_*basal*_ for WT in presence of the induced feedback loop. The PE on x-axis corresponds to estimated value of the basal pathway activity (*k*_1_) while The PE on y-axis corresponds to estimated value of the basal deactivation rate (*P*_*basal*_).

### Estimated parameters

1. *k*_1_ = 1078.4 nM^−1^ min^−1^
2. *S*_*basal*_ = 86.68 min^−1^
3. *P*_*basal*_ = 2036.6 min^−1^
4. *j*_*x*_ = 0.7653 nM min^−1^
5. 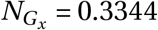
6. *d*_*x*_ = 0.0304 min^−1^
7. *K*_*GFP*_ = 3.7001 nM
8. *j*_*GFP*_ = 0.0594 nM min^−1^
9. *d*_*GFP*_ = 0.0363 min^−1^
10. 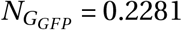
11. *k*_2_ = 2652.9 nM^−1^ min^−1^
12. *j*_*P*_ = 0.0174 nM min^−1^
13. 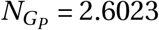
14. *K*_2_ = 0.2081 nM
15. *d*_*P*_ = 0.0174 min^−1^

